# Ivermectin Reduces Withdrawal-Induced Alcohol Intake via CeA GABAergic Enhancement

**DOI:** 10.1101/2025.10.14.682432

**Authors:** Paola Campo, Ran Qiao, Michelle R. Doyle, Daniel Munro, Benjamin J. Johnson, Abraham A. Palmer, Marsida Kallupi, Giordano de Guglielmo

## Abstract

Although FDA-approved medications for alcohol use disorder (AUD) are available, their efficacy varies across patients, highlighting the need for novel therapeutics that address inter-individual differences in disease etiology and treatment response. Genetic models, particularly heterogeneous stock (HS) rats, recapitulate human-like genetic diversity and behavioral heterogeneity, enabling the dissection of individual differences in vulnerability to AUD and pharmacotherapeutic sensitivity.

P2X4 receptors, which are encoded by the gene *P2rx4*, are ATP-gated ion channels inhibited by ethanol and abundantly expressed in neurons found in reward and stress circuits. P2X4 receptors have emerged as key modulators of ethanol sensitivity and consumption in preclinical models. Here, we genetically predicted *P2rx4* expression in whole brain in a cohort of 130 HS rats exposed to chronic intermittent ethanol (CIE) vapor and phenotyped for self-administration during acute abstinence. Rats were dichotomized into high- and low-predicted expression groups. Higher predicted *P2rx4* expression was associated with increased post-vapor intake and escalation. In 32 CIE-escalated rats, ivermectin, a positive allosteric modulator of P2X4 receptors, dose-dependently reduced drinking. We stratified rats into three groups: non-responders, mild responders, and high responders. Electrophysiological recordings from CeA slices revealed that ivermectin differentially enhanced GABAergic IPSCs: high-responders exhibited sustained increases in IPSC frequency and selective amplitude reductions, while the two other groups showed transient frequency increases. All groups displayed prolonged rise times, however non-responders showed extended decay times. These findings suggest that *P2rx4* upregulation serves as a vulnerability marker for dependence-like behaviors, with ivermectin attenuating withdrawal-driven alcohol consumption by enhancing CeA GABAergic inhibition.

## 1. Introduction

Alcohol use disorder (AUD) is a chronic relapsing condition marked by compulsive drinking, loss of control over intake, and negative emotional withdrawal states that drive relapse through negative reinforcement (Koob and Le Moal, 1997). With lifetime prevalence rates exceeding 29% in the United States and annual economic costs surpassing $249 billion, AUD contributes significantly to morbidity and mortality (Sacks et al., 2015). Current FDA-approved medications (disulfiram, naltrexone, acamprosate) have demonstrated efficacy in reducing drinking in clinical trials but are only effective in a subset of patients (Hartwell and Kranzler, 2019). Fewer than one in five people with AUD receive any treatment, and only a small fraction of those are prescribed medications, reflecting limitations due to genetic, neurobiological and environmental heterogeneity among patients (Biernacka et al., 2021; Hartwell and Kranzler, 2019; Mason and Heyser, 2021). These observations underscore the need for new therapeutic targets and a precision-medicine approach that accounts for individual differences.

Preclinical models that capture genetic diversity are therefore essential. The N/NIH outbred heterogeneous stock (HS) rats, derived from eight inbred strains (Solberg Woods and Palmer, 2019) displays broad phenotypic variation in ethanol drinking and related behaviors (Doyle et al., 2025).

In recent years, we and others have phenotyped thousands of HS rats for addiction-related traits, creating powerful datasets that capture behavioral variability across a genetically diverse population (Carrette et al., 2021; de Guglielmo et al., 2024; Doyle et al., 2025; Gunturkun et al., 2022; King et al., 2025; Kuhn et al., 2025b; Lara et al., 2024). This large-scale effort enables genome-wide association studies (GWAS) (Kuhn et al., 2025a; Solberg Woods and Palmer, 2019) and predictive modeling of gene expression (Zhou et al., 2023). By integrating genotype data with brain tissue expression data (www.ratgtex.org; Munro et al 2022), we can estimate genetically regulated expression (GReX) of specific genes across individuals (Ehsan et al., 2024; Gusev et al., 2016). This approach allows us to functionally prioritize candidate genes and stratify HS rats based on predicted expression levels, facilitating investigations into how genetically driven variation in the expression of promising targets influences addiction-like behaviors.

One promising target is the purinergic P2X4 receptor (P2X4R), encoded by the gene *P2rx4*, a ligand-gated ion channel highly expressed in the central nervous system, where it facilitates ATP-mediated calcium influx and modulates GABAergic, glutamatergic, glycinergic, and cholinergic neurotransmission (Coddou et al., 2011). Enriched in mesolimbic structures like the ventral tegmental area (VTA) and nucleus accumbens (NAc), P2X4Rs influence reward processing and reinforcement (Pierce and Kumaresan, 2006; Xiao et al., 2008). Acute ethanol desensitizes P2X4R function at intoxicating doses, (Davies et al., 2005; Ostrovskaya et al., 2011), whereas chronic exposure upregulates P2rx4, particularly in microglia, promoting neuroinflammation and BDNF-dependent adaptations, (Gofman et al., 2014).

Strain-dependent work links P2rx4 to alcohol intake: in recombinant inbred rats, whole-brain P2rx4 expression negatively correlates with voluntary ethanol consumption (Tabakoff et al., 2009), but in high-alcohol-drinking (HAD) rats lentiviral P2rx4 knockdown in the posterior VTA decreases drinking (Franklin et al., 2015).

Ivermectin (IVM), an antiparasitic agent used in veterinary and clinical medicine (Gonzalez et al., 2012), is a well-characterized positive allosteric modulator of P2X4Rs that antagonizes ethanol’s inhibitory effect on the receptor (Asatryan et al., 2014). In rodent studies, IVM significantly reduces voluntary ethanol consumption and self-administration (Asatryan et al., 2014; Yardley et al., 2012). Importantly, IVM’s alcohol-reducing effects are attenuated in P2X4-knockout mice, indicating that its action is at least partly P2X4-dependent (Khoja et al., 2018). Building on these findings, the present study harnesses the genetic diversity of HS rats to dissect *P2rx4’s* contributions to alcohol dependence phenotypes. Employing predictive models from RatGTEx, we imputed genetically regulated whole-brain *P2rx4* expression (GReX) across phenotyped individuals and dichotomized them into high- and low-predicted cohorts. We hypothesized that elevated predicted *P2rx4* expression would be associated with an exacerbated escalation of ethanol self-administration following chronic intermittent ethanol vapor (CIE) exposure, a paradigm that induces dependence-like neuroadaptations via enhanced negative reinforcement and withdrawal hypersensitivity (Doyle et al., 2025).

To evaluate therapeutic implications, we assessed ivermectin’s capacity to mitigate withdrawal-associated drinking in CIE-escalated rats. The central amygdala (CeA) plays an important role in mediating negative affective states during alcohol dependence through dysregulated GABAergic, and glutamatergic signaling (Gilpin and Roberto, 2012). To investigate this, we performed *ex vivo* slice electrophysiology in CeA neurons to quantify ivermectin’s modulation of GABAergic inhibitory postsynaptic currents (IPSCs) and correlate individual sensitivity with baseline synaptic parameters. This approach aimed to elucidate P2X4R-mediated mechanisms in CeA circuitry underlying vulnerability to AUD and inform precision pharmacotherapeutic strategies.

## 2. Materials and Methods

### 2.1 Subjects

Heterogeneous stock (HS) rats (N=163, 82 males and 81 females) were provided by The Center for Genetics, Genomics, and Epigenetics of Substance Use Disorders in Outbred Rats (P30DA060810; HS West: McwiWfsmAap:HS <u>#155269102</u>, RID:RGD_155269102). At weaning, rats were implanted with radio-frequency identification (RFID) chips for individual identification. Before every experiment, the RFID chip was scanned to confirm the rat’s identity, ensuring accurate data linkage and high-fidelity results.

Rats were pair-housed under a 12hour/12hour reverse light-dark cycle (lights on at 20:00 during the non-dependent phase or 21:00 during the dependent phase) in temperature-controlled (20– 22°C) and humidity-controlled (45–55%) rooms. They had *ad libitum* access to tap water and chow (Envigo Teklad Rat Food Diet 8604) outside of experimental sessions. With few exceptions (e.g., training sessions), experiments were conducted between 10:00 – 13:00, corresponding to 1-5 hours into the dark cycle. Rats were handled at least three times before initiating experimental procedures. All the procedures were performed in accordance with the ARRIVE guidelines (Kilkenny et al., 2010) adhered National Research Council’s Guide for the Care and Use of Laboratory Animals (National Research Council, 2011), and were approved by the Institutional Animal Care and Use Committee of the University of California San Diego.

### 2.2 Drugs

Ethanol (95%) was diluted to 10% (v/v) with purified tap water and delivered at a volume of 0.1 mL per reinforcer during operant self-administration sessions. Ivermectin (MedChemExpress, Monmouth Junction, NJ, USA) was freshly dissolved in 95% physiological saline (0.9% NaCl) and 5% Tween 80, and administered intraperitoneally (i.p.) 4 hours before behavioral testing at doses of 0, 1, 2.5, 5, and 10 mg/kg body weight, selected to span a range encompassing previously reported efficacious doses for reducing ethanol intake in rodent models (Yardley et al., 2012). For ex vivo slice electrophysiology studies in the central amygdala, ivermectin was bath-applied at concentrations of 0, 1, 5, and 10 μM, which have been shown to positively modulate P2X4 receptors and counteract ethanol’s inhibitory effects in heterologous expression systems and neuronal preparations (Khakh et al., 1999; Stokes et al., 2017).

### 2.3 Self-administration procedure

#### 2.3.1 Apparatus

Self-administration sessions were conducted in standard operant conditioning chambers (Med Associates, St. Albans, VT, USA). Two retractable levers were positioned on the right wall, with a sipper cup containing two wells situated between them. At the start of the session, both levers were extended into the chamber and remained accessible throughout the session. Two automated syringe drivers, each equipped with a 30-ml syringe, delivered ethanol and water solutions into the right and left wells, respectively.

#### 2.3.2 Training

To train rats in oral self-administration, they were placed into operant chambers for a 16-hr self-administration session, during which tap water (0.1 ml/reinforcer) was delivered to a sipper cup upon completion of a fixed ratio (FR) 1 schedule of reinforcement. Two days later, rats underwent another 16-hr self-administration session in the operant chambers, where 10% v/v ethanol in tap water (0.1 ml/reinforcer) was delivered via an FR1 schedule. During both sessions, chow was available *ad libitum,* but access to water or ethanol was contingent on active lever responses. Responses during the 0.6-sec timeout were recorded but had no programmed consequences.

#### 2.3.3 Ethanol-water choice pre-dependence

After initial training, rats were allowed to self-administer ethanol (10% v/v) (right lever) or water (left lever) on FR1:FR1 schedule during daily 30-min sessions, with each lever response delivering 0.1 ml of the respective solution.

#### 2.3.4 Chronic intermittent ethanol vapor exposure to induce dependence

Rats were rendered physically dependent using the chronic intermittent ethanol (CIE) vapor exposure model (de Guglielmo et al., 2023; Doyle et al., 2024; Kononoff et al., 2018; Vendruscolo and Roberts, 2014). Throughout the dependence phase, rats were pair-housed (two per cage) in ethanol vapor chambers (La Jolla Alcohol Research Inc, La Jolla, CA). Ethanol vapor was delivered for 14 hours daily, followed by a 10-hour off period, during which rats remained in the chambers. Rats were removed from the chambers only for self-administration sessions conducted during acute withdrawal (7–10 hours post-vapor exposure, see below). Over two weeks, ethanol vapor levels were gradually increased, and blood ethanol concentrations (BECs) were measured twice weekly until reaching an average of approximately 180 mg/dl, with most individual rats ranging between 150-225 mg/dl.

#### 2.3.5 Self-administration during ethanol dependence

Following induction of physical dependence via the chronic intermittent ethanol vapor exposure (CIE) model, rats resumed self-administration sessions on Mondays, Wednesdays, and Fridays during acute withdrawal (7-10 hours post vapor cessation).

The self-administration procedure for ethanol and water mirrored pre-dependence conditions, with the following modifications: 12 sessions were conducted to allow for escalation, followed by two quinine sessions and a progressive ration (PR) assessment.

#### 2.3.6 Blood alcohol Levels (BALs)

BALs were measured by collecting 0.1 ml of tail blood into a heparinized tube after pricking the tail vein with an 18G needle. Blood was spun in a centrifuge at 3000 RPMs for 13 minutes to separate plasma. Plasma was then analyzed using gas chromatography (de Guglielmo et al., 2023) or using an Analox AM1 Alcohol Analyser (Stourbridge, UK).

### 2.4 Calculating predicted gene expression using FUSION

Genetically predicted expression of *P2rx4* was predicted for the cohort of 130 HS rats phenotyped for alcohol dependence-like behaviors that were part of the UCSD Alcohol BioBank (Doyle et al., 2025). Genetic prediction was based on cis-regulatory expression models derived from bulk RNA-sequencing data from whole-brain hemisphere tissue (n ≈ 340 samples). Gene expression was quantified as transcript abundances using kallisto, (Bray et al., 2016), and then inverse-normal transformed. Twenty principal components from gene expression and five principal components from genotypes were used as covariates, following the Pantry framework (Munro et al., 2024). An elastic net predictive model for *P2rx4* was generated from gene expression, genotypes, and covariates using FUSION (Gusev et al., 2016), converted to individual-level predictors using FUSION’s make_score.R script, and applied to genotypes (VCF format) of the 130 HS rats via PLINK2 (Chang et al., 2015) with the --score parameter, yielding normalized values as Z-scores representing relative deviation from the cohort mean.

### 2.5 Central Amygdala (CeA) Slice Preparation for Electrophysiology

HS rats were anesthetized with 3–5% isoflurane during acute withdrawal (dependent rats) or at the same time of day (ethanol-naïve rats) using. Brains were rapidly removed, and 300 μm transverse slices were cut (de Guglielmo et al., 2019; Kallupi et al., 2020) using a Vibratome Series VT1000S (Technical Products International, St. Louis, MO, USA). Slices were incubated in an interface configuration with artificial cerebrospinal fluid (aCSF; 130 mM NaCl, 3.5 mM KCl, 1.25 mM NaH₂PO₄, 1.5 mM MgSO₄·7H₂O, 2.0 mM CaCl₂, 24 mM NaHCO₃, 10 mM glucose) warmed to 31°C for ∼30 min. After this incubation period, the slices were completely submerged and continuously superfused (3 mL/min) with room temperature aCSF, equilibrated with 95% O₂/5% CO₂ throughout the experiment.

#### 2.5.1 Whole-Cell Recording of Spontaneous Inhibitory Postsynaptic Currents (sIPSCs)

We recorded from neurons in the medial CeA that were visualized in brain slices using infrared differential interference contrast optics and CCD cameras (Exi Aqua and Q imaging). A 60× water immersion objective (Olympus) was used to identify and approach CeA neurons. Whole-cell voltage-clamp recordings were performed using a Multiclamp 700B amplifier (Molecular Devices), low-pass filtered at 2–5 kHz, digitized (Digidata 1440A; Molecular Devices), and stored on a computer using pClamp 10 software (Axon Instruments). All voltage-clamp recordings were conducted in gap-free acquisition mode with a sampling rate of 10 kHz. Patch pipettes (4–6 MΩ) were pulled from borosilicate glass (King Precision) and filled with an internal solution (70 mM KMeSO₄, 55 mM KCl, 10 mM NaCl, 10 mM HEPES, 2 mM MgCl₂, 2 mM Na-ATP, and 0.2 mM Na-GTP). Ivermectin (0, 1, 5, or 10 μM) and bicuculline (30 μM; to confirm GABAergic mediation) were constituted in aCSF and applied by bath superfusion (Kallupi et al., 2014b; Kallupi et al., 2013).

Spontaneous inhibitory postsynaptic currents (sIPSCs) were recorded while superfusing the slices with glutamate receptor blockers 6-cyano-7-nitroquinoxaline-2,3-dione (DNQX; 20 μM) and DL-2-amino-5-phosphonovalerate (DL-AP5; 30 μM). All cells were clamped at –60 mV for the duration of the recording.

The blockers were applied for at least 6 min before any other manipulation, followed by a baseline recording period of 6–8 min (with only the last 3 min used for analysis) (Kallupi et al., 2014a). Ivermectin was then bath-applied for 12–15 min to assess its effects, with the entire period analyzed in 3-min intervals. This was succeeded by an 8–10 min washout period (last 3 min analyzed). Finally, bicuculline was bath-applied for 3–6 min to verify the GABAergic nature of the sIPSCs (full period analyzed). Cells were excluded if membrane potential was outside the range of 60–65 mV or if series resistance changed by more than 20%.

### 2.6 Statistical Analysis

Statistical analyses of behavioral and electrophysiological data were performed using GraphPad Prism (version 10.0; GraphPad Software, Boston, MA, USA). Operant self-administration trajectories were evaluated using two-way repeated-measures ANOVA for time × predicted expression interactions, with separate pre- and post-vapor analyses to isolate phase effects; Bonferroni’s multiple comparisons probed session-specific escalations from the last pre-vapor baseline. Escalation in the ivermectin cohort employed one-way repeated-measures ANOVA with Bonferroni’s post-hocs, while averaged pre- versus post-vapor intake and group differences in terminal metrics were assessed via paired or unpaired two-tailed t-tests, respectively. Blood alcohol levels were analyzed by repeated-measures ANOVA with Bonferroni’s corrections. Ivermectin dose-responses utilized repeated-measures ANOVA for treatment effects, with two-way extensions for sex interactions and stratified analyses; Bonferroni’s post-hocs identified dose-specific changes. Responsivity stratification Z-scores were compared via one-way ANOVA with Bonferroni’s tests; correlations used Pearson’s r for linear regression. Electrophysiological parameters were normalized to baseline (100%) and evaluated with one-sample t-tests versus 100%; temporal dynamics involved two-way repeated-measures ANOVA with Bonferroni’s post-hocs for group differences, and within-group one-sample t-tests assessed bin-specific changes. Significance was set at P < 0.05. Data are presented as mean ± SEM.

## 3. Results

### 3.1 Genetically Regulated P2rx4 Expression and Escalation of Ethanol Self-Administration Following Chronic Intermittent Exposure

HS rats phenotyped for the UCSD Alcohol Biobank (Doyle et al., 2025) were stratified into low-predicted (n = 77) and high-predicted (n = 54) *P2rx4* expression groups based on imputed whole-brain genetically predicted expression (GReX) Z-scores, derived from a model with a cis-heritability (h²) of 0.61.

Figure 1A depicts the number of rewards earned for 10% (v/v) ethanol across pre-vapor baseline sessions and post-vapor sessions following chronic intermittent ethanol (CIE) vapor exposure. To evaluate overall trajectories and escalation, a two-way repeated-measures ANOVA was conducted across all sessions, revealing a significant time × predicted expression interaction (P = 0.0237), main effect of time (P < 0.0001), and main effect of predicted expression (P = 0.0351), demonstrating differential responding over time between groups.

**Figure 1.**
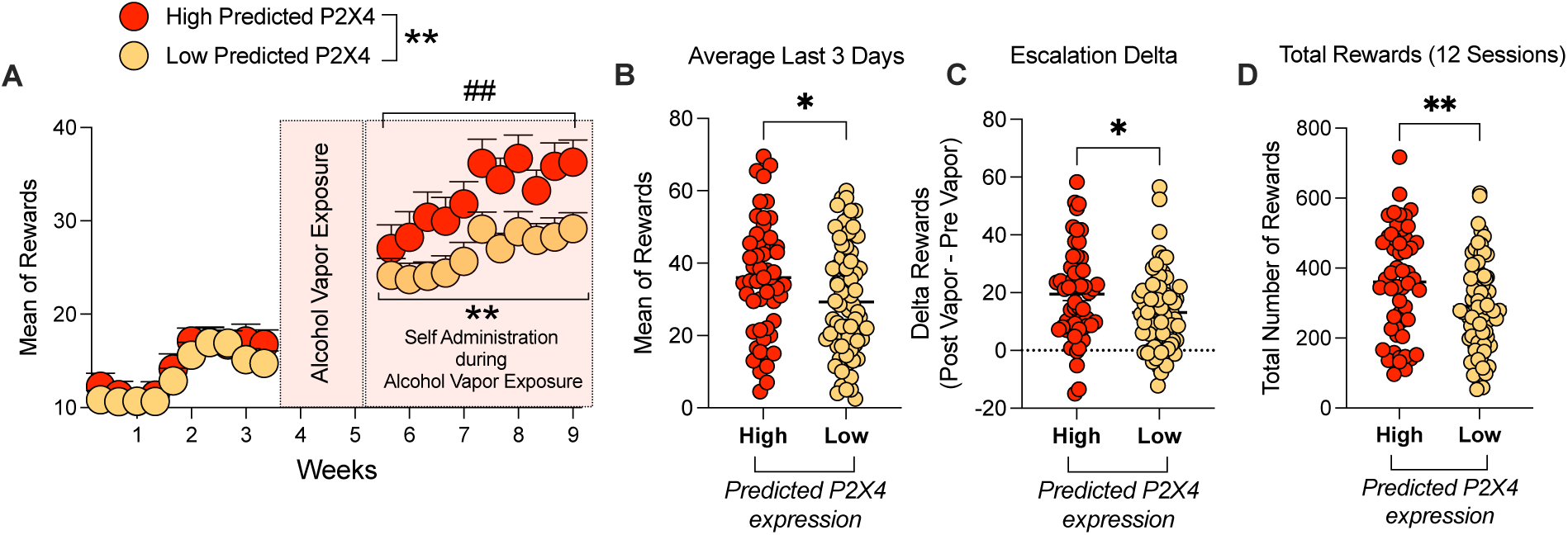
Genetically regulated P2rx4 expression predicts escalation of ethanol self-administration in heterogeneous stock (HS) rats following chronic intermittent ethanol (CIE) vapor exposure. (A) Temporal trajectory of operant responding, expressed as the number of rewards earned for 10% (v/v) ethanol (0.1 mL/reinforcer), across pre-vapor baseline sessions (left) and post-vapor escalation sessions (right). Data are presented as mean ± SEM for low-predicted (n = 77; mustard circles) and high-predicted (n = 54; red circles) groups based on imputed whole-brain genetically predicted gene expression (GReX) of P2rx4. The overall two-way repeated-measures ANOVA revealed a significant time × prediction interaction (P = 0.0237). Bonferroni’s post-hoc tests indicated significant differences from the last pre-vapor session (session 10) to each post-vapor session in both groups (all P < 0.01; # for low-predicted, * for high-predicted; ##/**P < 0.01; symbols placed above respective data points). Separate ANOVAs for pre- and post-CIE periods revealed no group differences pre-vapor but a significant main effect of predicted expression post-vapor (P = 0.0083, **P < 0.01). (B) Average rewards during the final three post-vapor sessions, showing elevated intake in the high-predicted group (unpaired t-test: P = 0.0179). (C) Escalation delta, calculated as the difference in rewards between the last three post-vapor and last three pre-vapor sessions, demonstrating greater escalation in the high-predicted group (P = 0.0146). (D) Cumulative rewards across all 12 post-vapor sessions, indicating higher total intake in the high-predicted group (P = 0.0049). Data in (B–D) are mean ± SEM; *P < 0.05, **P < 0.01 (unpaired two-tailed t-tests).

Bonferroni’s multiple comparisons tests, comparing the last pre-vapor session (session 10) to each of the 11 post-vapor sessions, confirmed significant escalation in both groups: for low-predicted rats, all comparisons were significant (P < 0.0001); for high-predicted rats, comparisons were significant with P values ranging from < 0.01 (session 17) to < 0.0001 (sessions 19–27), indicating robust post-vapor increases relative to pre-vapor baseline in both cohorts, with greater magnitude in the high *P2rx4* predicted group as reflected by the interaction. Complementing this, separate ANOVAs for pre- and post-vapor periods were performed to isolate phase effects. For the pre-vapor period, there was no significant time × prediction interaction (0.1% of variation, P = 0.7326) or main effect of predicted *P2rx4* expression (0.2% of variation, P = 0.5235), with only a main effect of time (4.9% of variation, P < 0.0001), demonstrating equivalent behavioral acquisition across groups. For the post-vapor period, there was no significant time × predicted *P2rx4* interaction (0.2% of variation, P = 0.7650), but main effects of time (2.3% of variation, P < 0.0001) and predicted *P2rx4* expression (3.0% of variation, P = 0.0083) were evident, demonstrating sustained group differences during dependence-like states.

Building on these analyses, we then examined terminal post-vapor performance, where the average number of rewards earned during the final three sessions (Figure 1B) was significantly higher in the high *P2rx4* predicted group compared to the low *P2rx4* predicted group (unpaired two-tailed t-test: t(129) = 2.399, P = 0.0179), demonstrating enhanced intake in rats predicted to have higher expression of *P2rx4*. To quantify escalation, we calculated the delta change in rewards (last three post-vapor minus last three pre-vapor sessions; Figure 1C), which was significantly greater in the high *P2rx4* predicted group (t(129) = 2.477, P = 0.0146), demonstrating amplified withdrawal-driven increases. Finally, cumulative responding across the 12 post-vapor sessions (Figure 1D) was significantly higher in the high-predicted group (t (129) = 2.861, P = 0.0049), demonstrating greater overall escalation burden associated with higher predicted P2rx4 expression.

### 3.2 Ivermectin Attenuates Escalated Ethanol Self-Administration in Dependent HS Rats

A subset of 32 heterogeneous stock (HS) rats (16 males, 16 females) underwent operant training for 10% (v/v) ethanol self-administration followed by chronic intermittent ethanol (CIE) vapor exposure to induce dependence, enabling assessment of escalation dynamics and pharmacological intervention with ivermectin. Figure 2A illustrates the temporal profile of ethanol intake (g/kg) across pre-vapor baseline sessions and post-vapor sessions, analyzed via one-way repeated-measures ANOVA, which revealed a significant main effect of time (F(7.746, 240.1) = 11.29, P < 0.0001), demonstrating progressive escalation post-exposure. Bonferroni’s multiple comparisons tests, comparing the last pre-vapor session (session 10) to each post-vapor session, confirmed significant increases in sessions 17, 18, 20, 21, and 22 (P < 0.01 to P < 0.001), with session 19 approaching significance (P = 0.059), demonstrating robust vapor-induced amplification of intake. Figure 2B depicts the average intake during the final three pre-vapor versus post-vapor sessions, where a paired two-tailed t-test indicated marked escalation (t (30) = 5.926, P < 0.0001), confirming that CIE reliably increases alcohol responding in this genetically diverse population. Concurrently, blood alcohol levels (BALs) were monitored to validate exposure intensity (Figure 2C), with repeated-measures ANOVA showing a significant effect of time (F(2.161) = 248.5, P < 0.0001); Bonferroni’s post-hocs revealed progressive elevations from week 5 to weeks 7-10 (P < 0.0001; vs. weeks 7–10), demonstrating titration to target levels (∼150–200 mg/dL).

**Figure 2.**
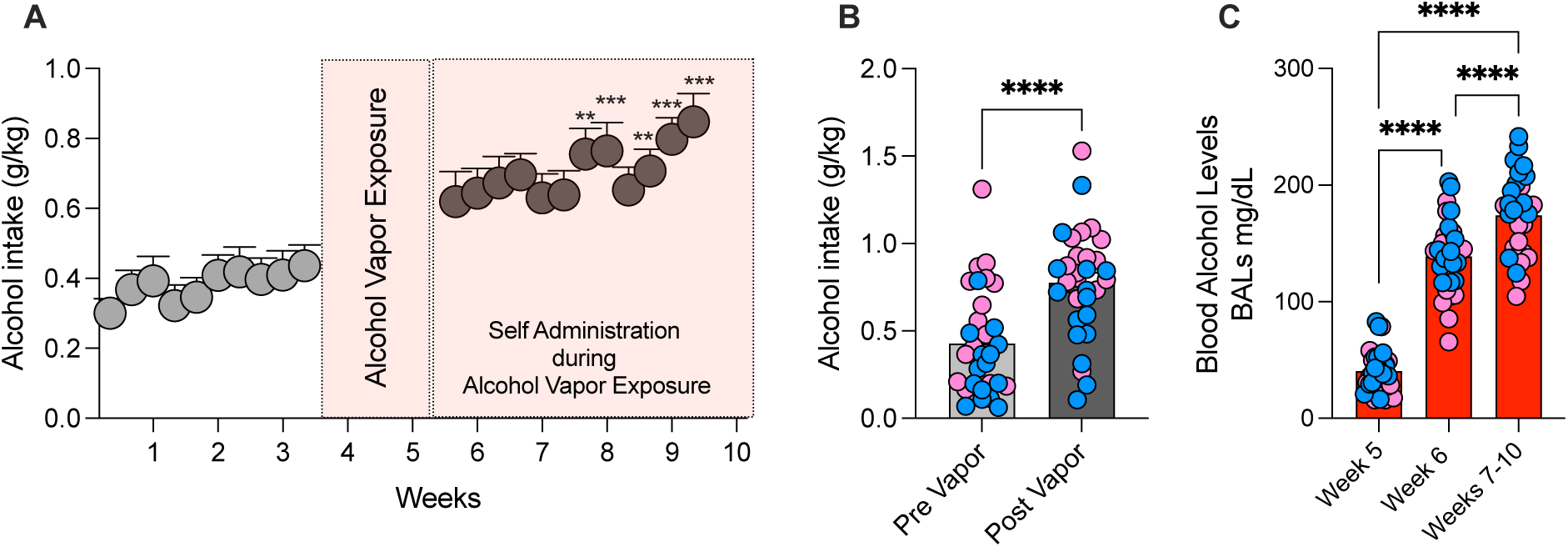
Escalation of ethanol self-administration following CIE vapor exposure in a subset of HS rats. (A) Temporal profile of operant ethanol intake (g/kg body weight) for 10% (v/v) ethanol (0.1 mL/reinforcer) across pre-vapor baseline sessions (left) and post-vapor escalation sessions (right, highlighted area). Data are presented as mean ± SEM for the cohort (n = 32; 16 males, 16 females). One-way repeated-measures ANOVA revealed a significant main effect of time (P < 0.0001), with Bonferroni’s post-hoc tests indicating significant differences from the last pre-vapor session (session 10) to post-vapor sessions 17, 18, 20, 21, and 22 (**P < 0.01, ***P < 0.001). (B) Average intake during the final three pre-vapor versus post-vapor sessions, showing significant escalation (paired t-test: ****P < 0.0001). (C) Blood alcohol levels (BALs; mg/dL) across vapor exposure weeks, with repeated-measures ANOVA indicating a significant effect of time (**P = 0.0056). Bonferroni’s post-hoc tests revealed progressive increases from week 5 to week 6 and weeks 7–10 (****P < 0.0001). Data are mean ± SEM, with individual data points shown as blue dots (males) and pink dots (females).

In the same dependent cohort, the effects of systemic ivermectin (0, 1, 2.5, 5, 10 mg/kg, i.p., administered 6 h prior) on post-vapor self-administration were evaluated using a Latin square design. Ethanol intake (g/kg), shown in Figure 3A, was analyzed by repeated-measures ANOVA (F(3.228, 96.83) = 6.901, P = 0.0002). Bonferroni’s post-hoc tests revealed dose-dependent reductions in ethanol consumption at 5 mg/kg (P < 0.01) and 10 mg/kg (P < 0.01), demonstrating significant attenuation of escalated consumption. Similarly, Figure 3B presents the number of rewards, with ANOVA indicating a significant treatment effect (F(3.336, 100.1) = 3.985, P = 0.0078) and post-hoc tests confirming reductions at 5 mg/kg (P < 0.05) and 10 mg/kg (P < 0.05), demonstrating ivermectin’s impact on alcohol intake without altering non-specific operant behavior, as water rewards remained unaffected (Figure 3C; F(2.216, 66.49) = 0.5922, P = 0.5728). To probe sex differences, Figure 3D displays intake separated by sex, analyzed via two-way repeated-measures ANOVA, which revealed no dose × sex interaction (F(3.167, 91.85) = 0.6823, P = 0.5728) but main effects of dose (F(3.167, 91.85) = 7.400, P = 0.0001) and sex (F(1, 29) = 13.05, P = 0.0011), demonstrating overall higher baseline intake in females and prompting sex-stratified analyses. For males (Figure 3E), ANOVA showed a significant treatment effect (F (2.847, 39.86) = 4.730, P = 0.0073), with post-hoc tests indicating reductions at 5 mg/kg (mean diff. vs. vehicle: 0.2099, P < 0.05) and 10 mg/kg (0.2485, P < 0.05), demonstrating comparable efficacy to the full cohort. For females (Figure 3F), a significant treatment effect emerged (F (2.760, 41.40) = 3.457, P = 0.0279), though post-hoc analysis revealed significance only at 10 mg/kg (mean diff. vs. vehicle: 0.2896, P < 0.05), demonstrating a potentially higher threshold for ivermectin’s anti-escalation effects in females.

**Figure 3.**
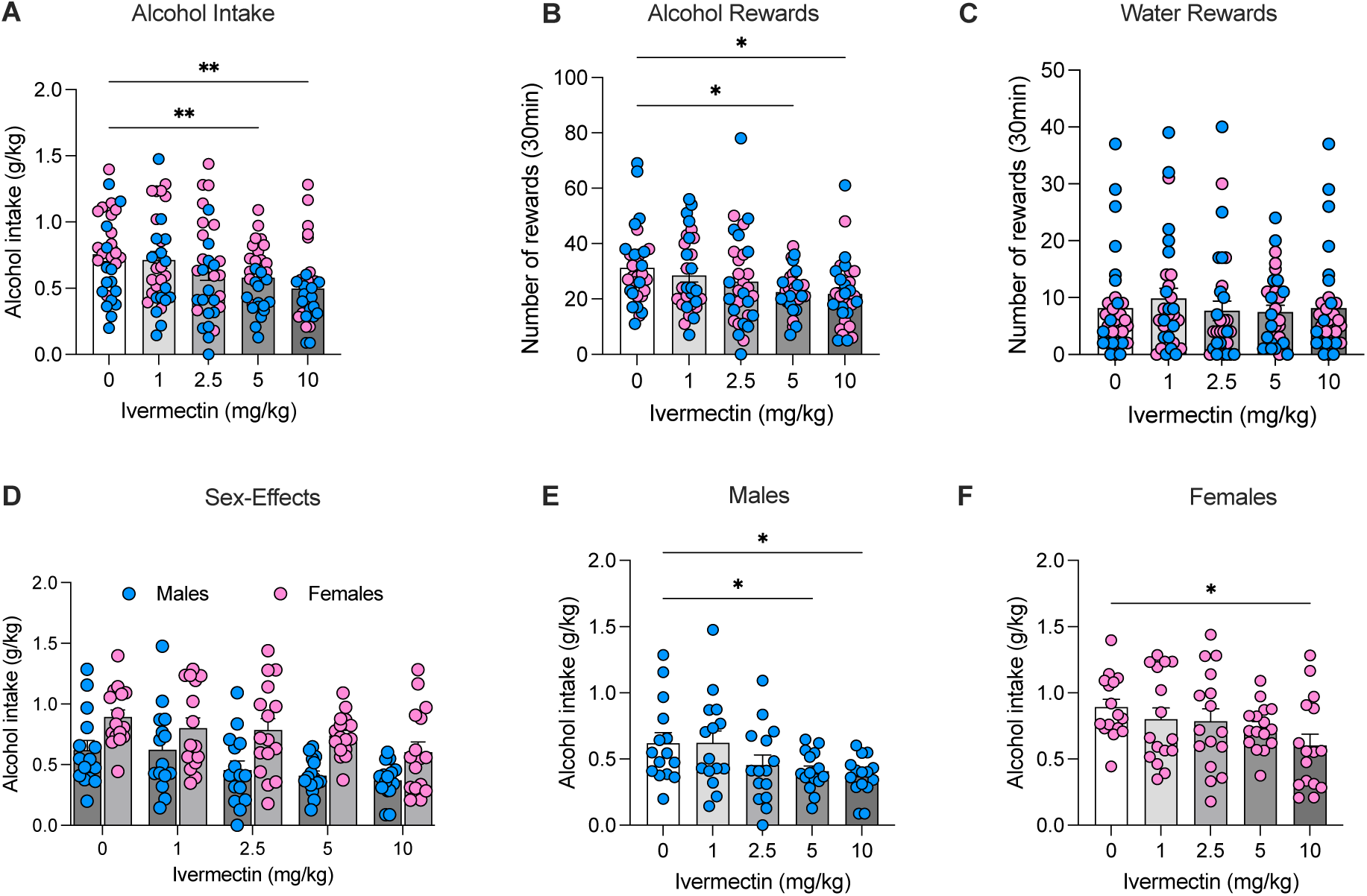
Dose-dependent effects of systemic ivermectin on escalated ethanol self-administration in dependent HS rats. All panels depict data from the same cohort (n = 32; 16 males, 16 females) tested in a Latin square design during post-vapor sessions. (A) Ethanol intake (g/kg body weight) across ivermectin doses (0, 1, 2.5, 5, 10 mg/kg, i.p., administered 6 h prior), with repeated-measures ANOVA showing a significant treatment effect (P = 0.0002). Bonferroni’s post-hoc tests indicated reductions versus vehicle at 5 and 10 mg/kg (**P < 0.01). Individual data points are shown as blue dots (males) and pink dots (females). (B) Number of rewards earned, with ANOVA revealing a significant treatment effect (P = 0.0078) and post-hoc tests showing reductions at 5 and 10 mg/kg (*P < 0.05). (C) Water rewards (non-specific operant control), unaffected by treatment (ANOVA: P = 0.5728). (D) Ethanol intake separated by sex (males: blue circles; females: pink circles), analyzed via two-way repeated-measures ANOVA, with main effects of dose (P = 0.0001) and sex (P = 0.0011) but no interaction (P = 0.5728). (E) Ethanol intake in males only (blue circles), with ANOVA indicating a significant treatment effect (P = 0.0073) and post-hoc tests showing reductions at 5 and 10 mg/kg (*P < 0.05). (F) Ethanol intake in females only (pink circles), with ANOVA showing a significant treatment effect (P = 0.0279) and post-hoc tests revealing reduction at 10 mg/kg (*P < 0.05). Data are expressed as mean ± SEM.

### 3.3 Stratification of Ivermectin Responsivity in Dependent HS Rats

To further dissect individual variability in ivermectin efficacy, the 32 dependent heterogeneous stock (HS) rats were stratified into non-responders (n = 11), mild responders (n = 10), and responders (n = 11) based on averaged Z-scores of delta decreases in ethanol intake across doses (1, 2.5, 5, 10 mg/kg), where positive Z-scores indicated greater responsivity. Figure 4A presents these averaged responsivity Z-scores by group, analyzed via one-way ANOVA, which revealed a significant group effect (F(2, 28) = 27.50, P < 0.0001); Bonferroni’s multiple comparisons tests confirmed differences between non-responders and responders (P < 0.0001) and mild responders and responders (P = 0.0003), but not between non-responders and mild responders (P = 0.0684), demonstrating graded levels of pharmacological sensitivity. Figure 4B illustrates a positive correlation between baseline post-vapor ethanol intake Z-scores and responsivity Z-scores (linear regression: slope = 0.5774 ± 0.1516, R² = 0.3334, P = 0.0007), demonstrating that rats with higher escalated intake exhibit greater reductions following ivermectin treatment. Blood alcohol levels (BALs) during vapor exposure did not differ across groups (Figure 4C; one-way ANOVA: F (2, 28) = 0.8313, P = 0.4460), demonstrating that responsivity stratification was independent of exposure intensity.

**Figure 4.**
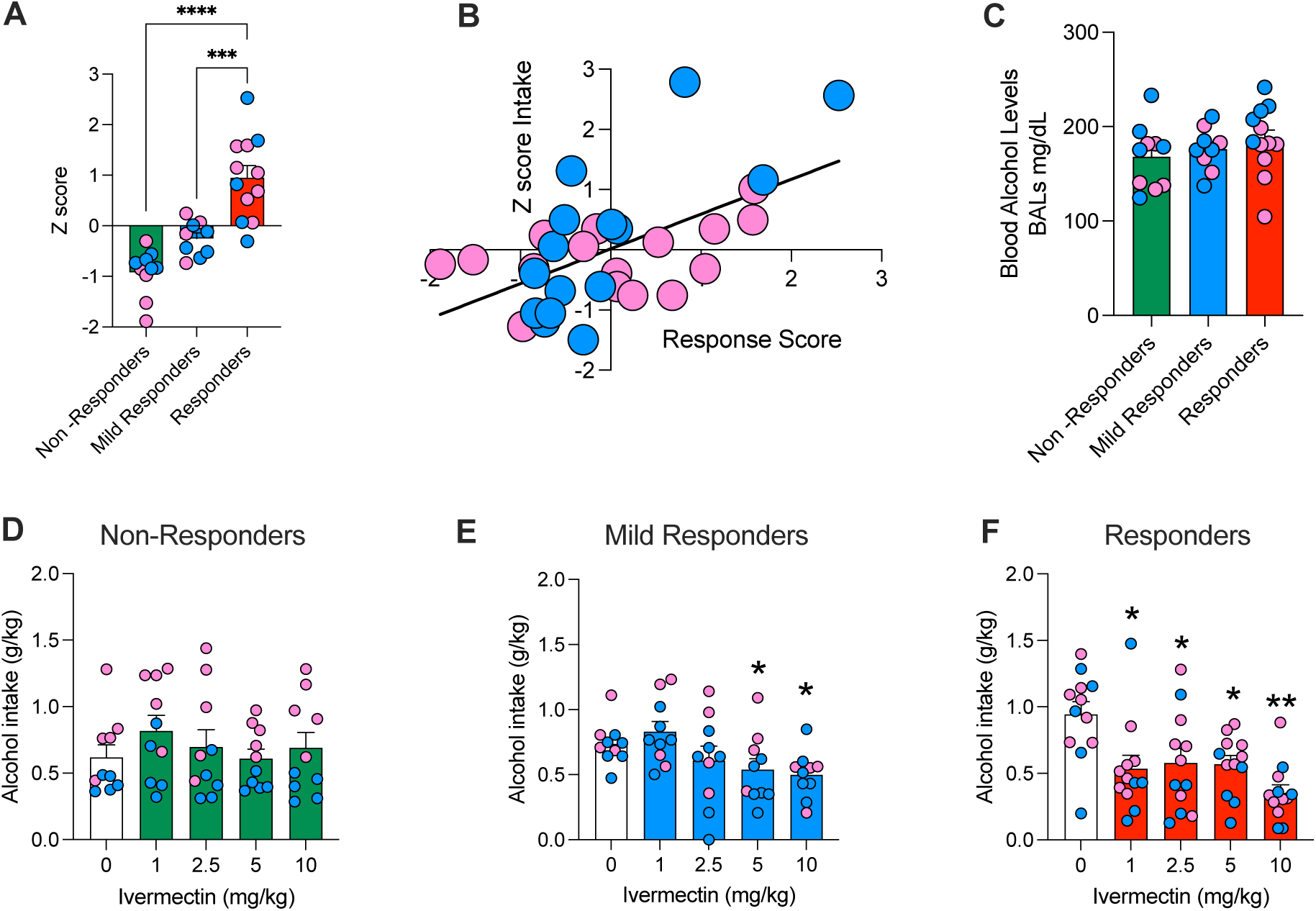
Stratification of ivermectin responsivity and its correlation with escalated ethanol intake in dependent HS rats. All panels depict data from the cohort (n = 32; 16 males, 16 females) stratified into non-responders (n = 11), mild responders (n = 10), and responders (n = 11) based on averaged Z-scores of deltas decrease in intake across ivermectin doses. (A) Averaged responsivity Z-scores by group, with one-way ANOVA showing a significant group effect (P < 0.0001). Bonferroni’s post-hoc tests indicated differences between non-responders and responders (****P < 0.0001) and mild responders and responders (***P = 0.0003). Data are presented as mean ± SEM. (B) Correlation between baseline post-vapor ethanol intake Z-scores and responsivity Z-scores (linear regression: R² = 0.3334, P = 0.0007). (C) Average blood alcohol levels (BALs; mg/dL) during vapor exposure showed no group differences (ANOVA: P = 0.4460). Data are presented as mean ± SEM. (D) Ethanol intake (g/kg) across doses in non-responders showed no significant treatment effect (ANOVA: P = 0.0774). Data presented as mean ± SEM. (E) Ethanol intake in mild responders exhibited a significant treatment effect (ANOVA: P = 0.0091) and post-hoc tests showing reductions at 5 and 10 mg/kg (*P < 0.05). Data are presented as mean ± SEM. (F) Ethanol intake in responders demonstrated a significant treatment effect (ANOVA: P < 0.0001) and post-hoc tests indicating reductions at all doses (*P < 0.05, **P < 0.01, ***P < 0.001). Data are presented as mean ± SEM. In all panels, individual data points are shown as blue dots (males) and pink dots (females).

Dose-response profiles were then examined within each stratum. For non-responders (Figure 4D), repeated-measures ANOVA showed no significant treatment effect (F (2.868, 25.81) = 2.582, P = 0.0774), demonstrating lack of ivermectin-induced attenuation of alcohol intake. In mild responders (Figure 4E), a significant treatment effect was observed (F (2.050, 18.45) = 6.066, P = 0.0091). Bonferroni’s post-hoc tests revealed reductions in ethanol consumption compared to vehicle at 5 mg/kg (P = 0.0253) and 10 mg/kg (P = 0.0416), demonstrating moderate sensitivity at higher doses. In responders (Figure 4F), ANOVA identified a robust treatment effect (F (3.230, 35.53) = 12.17, P < 0.0001), with post-hoc tests showing significant reductions at 1 mg/kg (P = 0.0073), 2.5 mg/kg (P = 0.0129), 5 mg/kg (P = 0.0048), and 10 mg/kg (P = 0.0003), demonstrating broad dose-dependent efficacy in this subgroup.

### 3.4 Ivermectin Modulates GABAergic Transmission in CeA Neurons from Dependent HS Rats Stratified by Behavioral Responsivity

Ex vivo whole-cell voltage-clamp recordings of spontaneous inhibitory postsynaptic currents (sIPSCs) in medial central amygdala (CeA) neurons were conducted to assess ivermectin’s effects on GABAergic transmission, comparing behavioral responders and non-responders (as defined in Figure 4) across doses (1, 5, and 10 μM; bath-applied for 12 min, with values averaged over this period and normalized to baseline 100%). One-sample t-tests versus 100% revealed dose-dependent enhancements in GABAergic transmission. Figure 5A shows that sIPSC frequency increased significantly only in responders at 10 μM (t (7) = 2.805, P = 0.0263), with no effect in non-responders (t (7) = 1.099, P = 0.3080). In contrast, sIPSC amplitude (Figure 5B) decreased significantly in responders at 10 μM (t (7) = 4.346, P = 0.0034), but not in non-responders (t (7) = 1.722, P = 0.1288). Rise time (Figure 5C) was significantly increased in non-responders at all doses tested (1 μM: t (5) = 5.073, P = 0.0039; 5 μM: t (7) = 2.792, P = 0.0268; 10 μM: t (7) = 2.653, P = 0.0328) and in responders at the 5 μM (t(6) = 3.268, P = 0.0171) and 10 μM doses (t(7) = 2.411, P = 0.0467), but not at the 1 μM dose (t(5) = 0.6394, P = 0.5507). Decay time (Figure 5D) increased only in non-responders at 10 μM (t (7) = 2.483, P = 0.0420), with no effects in responders (t (7) = 0.4738, P = 0.6501) or at lower doses.

**Figure 5.**
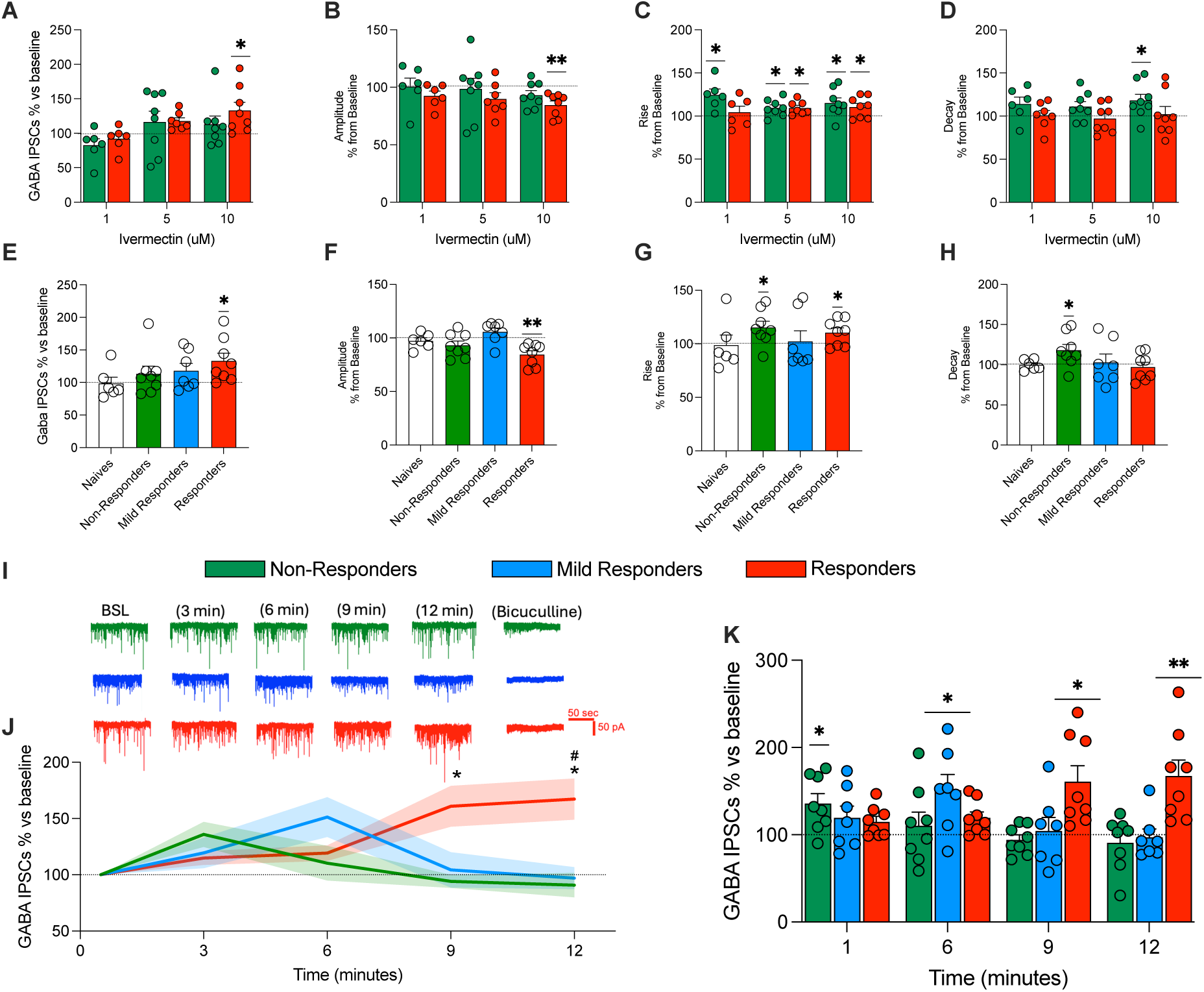
Ivermectin modulation of GABAergic sIPSCs in CeA neurons from dependent HS rats stratified by ivermectin responsivity. Dose-response effects (1, 5, 10 μM; averaged over 12 min) in responders versus non-responders: (A) Frequency, significantly increased only in responders at 10 μM (*P = 0.0263). (B) Amplitude, significantly decreased only in responders at 10 μM (**P = 0.0034). (C) Rise time, significantly increased in non-responders at all doses (**P < 0.05 to ***P < 0.01) and in responders at 5 μM and 10 μM (*P < 0.05). (D) Decay time, significantly increased only in non-responders at 10 μM (*P = 0.0420). Effects at 10 μM across all groups (ethanol-naïve controls, non-responders, mild responders, responders): (E) Frequency, significantly increased only in responders (*P = 0.0263). (F) Amplitude, significantly decreased only in responders (**P = 0.0034). (G) Rise time, significantly increased in non-responders (*P = 0.0328) and responders (*P = 0.0467). (H) Decay time, significantly increased only in non-responders (*P = 0.0420). Temporal dynamics at 10 μM: (I) Representative sIPSC traces shown for non-responders, mild responders, and responders. (J) Timeline of frequency changes (3-min bins), with two-way ANOVA showing a significant time × group interaction (P < 0.0001); Bonferroni’s post-hoc tests indicated differences at 9 minutes and 12 minutes (*P < 0.05 vs non-responders and # P < 0.05 vs mild responders). (K) Frequency per bin relative to baseline for each group, with significant elevations in non-responders at 3 minutes (*P = 0.0150), mild responders at 6 minutes (*P = 0.0281), and responders at 9 minutes (*P = 0.0120) and 12 minutes (**P = 0.0080). All data are presented as mean ± SEM (normalized to baseline [100%]); statistical analyses used one-sample t-tests against 100% unless otherwise noted.

Subsequent recordings at the maximally effective dose (10 μM) extended comparisons to include mild responders and ethanol-naïve controls. Figure 5E shows that sIPSC frequency was significantly increased only in responders (t (7) = 2.805, P = 0.0263), with no significant changes in ethanol-naive controls (t (5) = 0.1396, P = 0.8944), non-responders (t (7) = 1.099, P = 0.3080), or mild responders (t (6) = 1.606, P = 0.1595). Amplitude (Figure 5F) was significantly reduced only in responders (t (7) = 4.346, P = 0.0034), with no significant changes in ethanol-naïve controls (t (5) = 0.9278, P = 0.3961), non-responders (t (7) = 1.722, P = 0.1288), or mild responders (t (6) = 1.537, P = 0.1751). Rise time (Figure 5G) was significantly increased in non-responders (t (7) = 2.653, P = 0.0328) and responders (t (7) = 2.411, P = 0.0467), but not in ethanol-naïve controls (t (5) = 0.1396, P = 0.8944) or mild responders (t (6) = 0.2285, P = 0.8268). Decay time (Figure 5H) was significantly increased only in non-responders (t (7) = 2.483, P = 0.0420), with no significant effects observed in naïve controls (t (5) = 0.06264, P = 0.9525), mild responders (t (6) = 0.2803, P = 0.7887), or responders (t (7) = 0.4738, P = 0.6501). Temporal dynamics of sIPSC frequency during 10 μM ivermectin application were assessed in 3-min bins across non-responders, mild responders, and responders (Figure 5I; representative traces). A two-way repeated-measures ANOVA on the timeline (Figure 5J) revealed a significant time × group interaction (F (5.895, 58.95) = 7.167, P < 0.0001), main effect of time (F (2.948, 58.95) = 3.319, P = 0.0265), and a trend for group (F (2, 20) = 3.127, P = 0.0658). Bonferroni’s post-hoc tests revealed significant group differences at 9 minutes (non-responders vs. responders, P = 0.0219) and 12 minutes (non-responders vs. responders, P = 0.0122; mild responders vs. responders: mean difference = −70.25, P = 0.0195). Within-group comparisons to baseline (Figure 5K) showed transient elevations in sIPSC frequency: in non-responders at 3 minutes (t(7) = 3.201, P = 0.0150); in mild responders at 6 minutes (t (6) = 2.880, P = 0.0281); and in responders at 9 minutes (t (7) = 3.364, P = 0.0120) and 12 min (t(7) = 3.665, P = 0.0080).

## 4. Discussion

By integrating genetically predicted gene expression (GReX) of P2rx4 with behavior and central amygdala (CeA) physiology in heterogeneous stock (HS) rats, we identify a translationally tractable purinergic mechanism that (i) marks vulnerability to withdrawal-driven alcohol intake and (ii) can be pharmacologically leveraged with ivermectin (IVM), a positive allosteric modulator (PAM) of P2X4 receptors. HS rats with higher predicted *P2rx4* expression exhibited significant escalation after chronic intermittent ethanol (CIE) vapor, whereas systemic IVM attenuated post-vapor intake in a dose-dependent manner and produced distinctive signatures in CeA GABAergic synaptic transmission that tracked the behavioral response. Together, these results position *P2rx4* as a biologically informed stratifier and P2X4 receptor modulation as a precision-oriented avenue for AUD therapeutics.

Our behavioral–genetic findings align with convergent evidence that P2X4 receptors modulate ethanol-related phenotypes. At the receptor level, intoxicating ethanol acutely inhibits P2X4 receptors, whereas ivermectin (IVM) antagonizes this inhibition, establishing a bidirectional pharmacology relevant to drinking behavior (Asatryan et al., 2010). In vivo, reducing P2X4 function increases ethanol intake and attenuates IVM’s efficacy: in *P2rx4* knockout mice, baseline consumption is higher and the intake-lowering effect of IVM is ∼50% smaller than in wild type (Wyatt et al., 2014). Region-specific manipulations point the same way: posterior VTA *P2rx4* knockdown reduces intake in high-drinking rats, and ethanol directly antagonizes P2X4 receptors on VTA neurons ex vivo, linking mesolimbic P2X4 signaling to reinforcement mechanisms (Franklin et al., 2015). In our HS cohort, higher genetically predicted *P2rx4* expression was associated with greater escalation only after CIE exposure (with no pre-CIE differences), indicating that the P2X4–drinking relationship is state dependent and becomes unmasked during withdrawal. Directionality of the effects may depend on brain region and state: global loss removes P2X4 across circuits, producing a pro-drinking phenotype and blunting IVM’s on-target action, whereas higher expression in withdrawal-engaged areas (e.g., VTA and amygdalar circuits) can facilitate escalation. This interpretation is consistent with the VTA data above and with evidence that alcohol can increase *P2rx4* expression in microglia (Gofman et al., 2014), changes that may contribute to circuit adaptations during chronic exposure, though we did not test microglial mechanisms here. Crucially, our CeA physiology ties the behavioral stratification to an inhibitory synaptic signature: IVM produced sustained increases in sIPSC frequency selectively in responders, with group-specific kinetic changes and minimal effects in non-responders, implying heterogeneity in where and how P2X4Rs are engaged during dependence, a pattern fully compatible with the regional/state-dependent model above.

A key translational point is brain exposure at the doses we used. IVM is detectable in rodent brain after 5–10 mg/kg i.p.; brain concentrations peak on a delayed timescale (≈9 h in mice), and the time course of efficacy mirrors central exposure (Yardley et al., 2012). The 4-h pretreatment we used sits on the ascending portion of that central PK profile, providing a pharmacological rationale for the robust behavioral effects at 5 and 10 mg/kg. Although P-glycoprotein limits ivermectin’s CNS penetration in some contexts (Geyer et al., 2009), preclinical strategies such as pairing with complementary agents, such as dihydromyricetin, increase exposure and/or efficacy and further reduce drinking in mice (Silva et al., 2020).

In the CeA, ethanol acutely facilitates presynaptic GABA release, and dependence produces a persistent hyper-GABAergic state supported by stress–neuropeptide signaling (Bajo et al., 2008; Roberto et al., 2004). Notably, increased CeA GABA transmission does not uniformly promote alcohol intake; several anti-drinking interventions enhance inhibitory tone rather than suppress it, underscoring that the therapeutic relation between inhibition and drinking is circuit- and state-dependent, not monotonic. For example, the GLP-1 analogue semaglutide reduces alcohol intake and increases CeA sIPSC frequency in rats (Chuong et al., 2023); neuroactive steroids (ganaxolone, pregnenolone), which positively modulate GABA A receptors, reduce operant ethanol self-administration in alcohol-preferring rats (Besheer et al., 2010); and topiramate, which facilitates GABA A currents and antagonizes AMPA/kainate, reduces drinking in randomized trials and animal models of high alcohol consumption (Breslin et al., 2010; Paparrigopoulos et al., 2011). Within this framework, our slice physiology data show that responders to IVM exhibit a sustained increase in sIPSC frequency, whereas non-responders display prominent kinetic signatures (reduced mean amplitude with prolonged rise and decay) without a durable frequency increase. A mechanistic interpretation is that, in responders, IVM effectively couples P2X4 receptor signaling to presynaptic release at inhibitory terminals, elevating action-potential–dependent GABA output; in non-responders, the dominant effect is postsynaptic integration, yielding smaller, slower events without greater release. Because IVM can modulate GABA A and glycine receptors at micromolar concentrations (Krusek and Zemkova, 1994; Shan et al., 2001), we cannot exclude some contribution from these sites (Kirson et al., 2020); however, the group-selective frequency increase confined to responders argues that presynaptic mechanisms (plausibly P2X4R-linked) are necessary for the behavioral phenotype (Asatryan et al., 2010). Importantly, IVM is quantifiable in rodent brain after systemic dosing in the same range used here, confirming central exposure at 5–10 mg/kg (Yardley et al., 2012).

The sex-effects in our dataset refine and extend this framework. Females consumed more alcohol than males during post-vapor testing, yet males responded to lower IVM doses. These findings align with CeA physiology reports demonstrating sex differences in acute ethanol sensitivity and neuromodulatory control of GABA transmission (Kirson et al., 2021). This sex-by-dose dissociation has practical implications for dose selection and timing in future clinical translation. Methodologically, our use of genetic prediction leverages cis-eQTL information to stratify individuals into two groups which can then be compared for behavioral differences. This is conceptually similar to transcriptome-wide association studies, which use genetically predicted gene expression to identify correlations between gene expression and complex behavioral or other traits (Gamazon et al., 2015; Gusev et al., 2016; Zhu et al., 2016).

Translationally, our dosing converges with prior rodent studies showing that 1–10 mg/kg IVM reduces drinking across paradigms and strains (Kosten, 2011; Silva et al., 2021; Yardley et al., 2015; Yardley et al., 2012), and that genetic or pharmacological manipulations of P2X4Rs bidirectionally modulate intake (Asatryan et al., 2014; Wyatt et al., 2014). Human feasibility and safety have been preliminarily demonstrated (30 mg oral IVM with intravenous alcohol was well tolerated but not efficacious at that single dose/regimen), underscoring the need to optimize exposure and patient selection (Roche et al., 2016). Looking ahead, translation will benefit from PK/PD-guided dosing that aligns central exposure with the post-dose window of maximal efficacy observed in mice (Yardley et al., 2012). In parallel, approaches that increase CNS target engagement or act on convergent pathways merit testing, such as efflux modulation at P-gp and co-treatment with dihydromyricetin (DHM), both of which enhanced anti-drinking effects preclinically (Silva et al., 2021; Silva et al., 2020). Mechanistically, cell-type–specific perturbations of *P2rx4* within CeA microcircuits and their mesolimbic inputs should be used to dissect microglial versus interneuronal contributions and to establish necessity/sufficiency for the synaptic signatures we observe. These efforts would be complemented by the development of P2X4-selective, brain-penetrant positive allosteric modulators and pharmacodynamic readouts of CeA inhibitory tone, enabling early human studies in genetically stratified cohorts informed by *P2rx4* expression signatures.

In summary, genetically predicted *P2rx4* identified individuals prone to dependence-like escalation, and ivermectin, at brain-active doses, attenuated withdrawal-driven intake while rebalancing CeA GABAergic inhibition in a way that mirrors behavioral sensitivity. The alignment among genetics, systems pharmacology, and synaptic physiology supports P2X4R modulation as a credible, mechanism-anchored strategy for AUD and provides a rationale for trial designs that pre-specify sex and *P2rx4* expression as stratification factors when evaluating P2X4R-targeting agents.

## Acknowledgements/Funding

This work was supported by National Institute on Alcohol Abuse and Alcoholism [R01 AA030048 to GdG, R01AA029688 to AAP and T32 AA007456 to MRD], by the National Institute on Drug Abuse [P30DA060810 to AAP], and the Preclinical Addiction Research Consortium (PARC) at the University of California San Diego.

The authors declare no competing financial interests.

## Abbreviations

aCSF: artificial cerebrospinal fluid
AUD: Alcohol Use Disorder
BALs: Blood Alcohol Levels
BECs: Blood Ethanol Concentrations
BDNF: Brain-Derived Neurotrophic Factor
CeA: Central Amygdala
CIE: Chronic Intermittent Ethanol
eQTL: expression Quantitative Trait Loci
FR: Fixed Ratio
GABA: Gamma-Aminobutyric Acid
GReX: Genetically Regulated Expression
GWAS: Genome-Wide Association Studies
HAD: High-Alcohol-Drinking
HS: Heterogeneous Stock
IPSCs: Inhibitory Postsynaptic Currents
IVM: Ivermectin
NAc: Nucleus Accumbens
P2X4R: P2X4 Receptor
*P2rx4*: Purinergic Receptor P2X 4 Gene
PAM: Positive Allosteric Modulator
PR: Progressive Ratio
RFID: Radio-Frequency Identification
sIPSCs: Spontaneous Inhibitory Postsynaptic Currents
TWAS: Transcriptome-Wide Association Studies
VTA: Ventral Tegmental Area

